# Continual evolution through coupled fast and slow feedbacks

**DOI:** 10.1101/513416

**Authors:** Meike T. Wortel, Han Peters, Juan A. Bonachela, Nils Chr. Stenseth

## Abstract

The Red Queen Hypothesis, which suggests that continual evolution can result from solely biotic interactions, has been studied in macroevolutionary and microevolutionary contexts. While microevolutionary studies have described examples in which evolution does not cease, understanding which general conditions lead to continual evolution or to stasis remains a major challenge. In many cases, it is unclear which experimental features or model assumptions are necessary for the observed continual evolution to emerge, and whether the described behavior is robust to variations in the given setup. Here, we aim to find the minimal set of conditions under which continual evolution occurs. To this end, we present a theoretical framework that does not assume any specific functional form and, therefore, can be applied to a wide variety of systems. Our framework is also general enough to cast predictions about both monomorphic and polymorphic populations. We show that the combination of a fast positive and a slow negative feedback causes continual evolution to emerge even from the evolution of one single evolving trait, provided that the ecological timescale is sufficiently separated from the timescales of mutation and the negative feedback. Our approach and results thus contribute to a deeper understanding of the evolutionary dynamics resulting from biotic interactions.

## 1 Introduction

The evolutionary dynamics of a species in a complex ecosystem can be driven by the properties of the species, by the interaction with coexisting species and their environment, and/or by external factors. To better understand to what extent the emerging ecological and evolutionary dynamics are caused by the species or by (intra- or inter-specific) biotic interactions, it is essential to study systems in the absence of any abiotic drivers. Isolating a system from all abiotic factors may lead to a static adaptive landscape, where adaptation follows a path towards a peak in that landscape—reachable or not ^1,2^. However, since the major part of any individual’s environment is typically composed of other (evolving) species, any species’ environment will normally change even without external abiotic variation, both through ecological and evolutionary changes. Hence, the resulting adaptive landscape is expected to be dynamic, potentially leading to continual co-evolutionary dynamics, where traits are oscillating, escalating, or chasing each other over evolutionary time ^3^.

Continual evolutionary dynamics are evolutionary dynamics that do not result in stasis. This means that, although the evolutionary dynamics could be apparently stable for some period of time, eventually sustained fluctuations (i.e. either periodic or irregular changes over time) will materialize for genotypes and phenotypes over long timescales. These dynamics are directly related to Red Queen dynamics, although the latter typically refers to systems of at least two species, whereas the more general term “continual evolutionary dynamics” includes systems with a single species. The so-called Red Queen (RQ) dynamics ^4^ is a concept that has had a major influence on micro- and macroevolutionary theory. The analysis of macro-evolutionary models led to the conclusion that the emergence of either RQ dynamics or stasis depends upon the nature of the within-system biotic interactions ^5,6,7,8^. This body of work, however, was not able to translate into ecological terms what the conditions for the emergence of the RQ are, although recent research found that symmetric competitive interactions are more likely to lead to stasis ^9^. From the micro-evolutionary perspective, theoretical work has focused on mechanistic descriptions of specific examples where the RQ can emerge ^3^. Theoretical studies of micro-evolutionary RQ dynamics mostly use methods based on adaptive dynamics and quantitative genetics ^10,11,12,13,14,15^. The adaptive dynamics approach assumes that the ecological dynamics have reached an equilibrium before evolution can occur, and studies the attempts of invading this stationary state of the resident by mutants, defined as individuals with a value of the adaptive trait that is slightly deviating from that of the resident. These assumptions enable a rigorous theoretical analysis of the system. The quantitative genetics approach, on the other hand, does not require an ecological equilibrium, but also requires the timescale separation between evolutionary adaptation and ecological dynamics. If the rate of evolutionary change is very slow, the quantitative genetics approach becomes similar to adaptive dynamics. Both methods assume that adaptive traits evolve along a fitness gradient. Following these theoretical frameworks, studies focussing on predator-prey or host-parasite systems have been able to reach conclusions about the conditions that increase or decrease the chance of RQ dynamics in specific settings (e.g. fast adaptation is less likely to lead to RQ dynamics ^11^, and RQ dynamics require an intermediate harvesting efficiency of the prey ^10^).

Most of the work mentioned above constrains the evolving population to be monomorphic, and therefore there is no guarantee that the same examples will lead to RQ dynamics in a polymorphic setting (which includes coexistence of subpopulations with different trait values). A polymorphic trait distribution arises easily with asexual reproduction or traits that are determined by a few loci, but can also arise when assortive mating develops (see the discussion in Kisdi et al. ^16^). Hence, the conditions that lead to the emergence of the RQ, or, more generally, continual evolution, in polymorphic populations remain elusive. Since most evolution experiments are done with microorganisms, and those are prone to showing polymorphic populations, this knowledge gap can prevent linking theoretical results to experimental data. Here, we fill this gap by studying the mechanisms leading to continual evolution without constraining the distribution of phenotypes that may be present in the population, and therefore the emergent continual evolution dynamics do not rely on the assumption of a monomorphic population. Additionally, we allow for interactions with (possible abiotic parts of) the environment and for mutations of small and large effects, which can be expected in such populations. In a previous study, we showed that micro-evolutionary RQ dynamics are possible in a model of a polymorphic bacterial system due to species interactions that lead to a changing adaptive landscape ^17^. This study was however limited to a specific set of eco-evolutionary interactions. Similarly, most of the theoretical microevolutionary RQ studies use specific functional forms for their analysis, hampering the generalisation of the obtained results. Meta-analyses (such as the one by Abrams ^18^) can provide some more general insights, but since many of these studies use similar equations, the breadth of the conclusions that can be drawn from them are still limited. To obtain general results and a broad understanding of what ecological interactions can cause evolutionary patterns such as continual evolution, we need as general models as possible. Here, we aim at extending the theoretical understanding of the conditions that lead to either continual evolution or stasis. For this purpose, we use a general model with a reduced set of assumptions regarding the form of the model functions. We find that a system with slow and fast feedback interactions in a polymorphic setting exhibits continual evolutionary dynamics in the presence of timescale separation, regardless of the size or effect of mutations.

## 2 Model description and the emerging eco-evolutionary dynamics

### 2.1 General model description

We model a population with genetic variation using differential equations that allow for mutations that are not constrained to be infinitesimal changes in the trait value. In order to focus our argument, we use a general model of a population density distribution *u*(*x, t*) over the trait space *x* and the distribution of environmental factors (i.e. abiotic factors and non-evolving species), *φ*(*y, t*), over the environment space *y*:

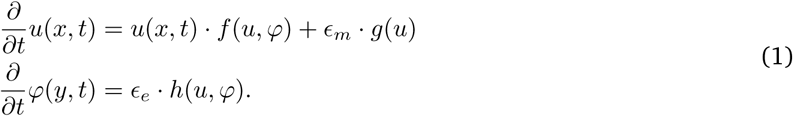

In this general case, the evolving trait is not specified and therefore its trait space could have any shape and dimensionality (i.e. any number of evolving traits). Distinct peaks for *u*(*x, t*) in the trait space, for example, represent a polymorphic trait distribution within a species if there can be exchange of individuals between these distinct peaks, or different species if there is no exchange between peaks (or very low exchange, in a macro-evolutionary setting). Similarly, the environment space, *y*, which consists of all non-evolving parts of the system, could have any form. The functional *f* = *f* (*x*) describes the growth of the population, which can depend on the population density distribution *u*(*x, t*) as well as on the environmental factors *φ*(*y, t*). The functional *g* describes the change in the population density due to mutations, which could reasonably be represented by a diffusion process ^9^. This change due to mutations is modeled in an unbiased manner, and the benefits of mutation result from the overall performance of the population. The functional *h* gives the rate of change in environmental factors, which can again depend on *u* and *φ*. Time scale differences between growth, environmental factors and mutations are captured in *ϵ*_*e*_ and *ϵ*_*m*_.

### 2.2 Continual evolutionary dynamics and the role of feedbacks

In this paper, we show the conditions that guarantee continual evolutionary dynamics in the generic model above, Eqs.(1). These conditions include a fast positive feedback and a slow negative feedback. The former is incorporated in the model through a positive effect of the trait density *u* on the population growth *f*. For the latter, the negative effect is implemented through how the dynamics of the environmental factors depends on the population density (i.e. how the functional *h* depends on *u*) and how environmental factors affect the growth of the population (i.e. how *φ* affects *f*); the slowness of the negative feedback is incorporated through the timescale variable *ϵ*_*e*_ ≪ 1.

Since non-evolving parts of the system are captured in the environmental factor *φ*, a biological example of the slow negative feedback could be a predator-prey interaction where the generation time of the predator is much slower than that of the prey, and where the predator targets preferentially a specific prey phenotype or prey type (e.g. bears preferring large salmon ^19^ or zooplankton preferring phytoplankton with a certain nitrogen:phosphorus ratio ^15^). The fast positive feedback is essentially an Allee effect on the population, as increases in the number of individuals with a certain trait value increases the associated growth rate. This could be due to, for example, cooperation in defense or feeding (e.g. when the evolving population is foraging on a certain vegetation area with sufficient resources, the more individuals forage in the same place, the better they are protected against predation). More examples of these feedbacks are listed in SI Table S3.

## 3 Results

### 3.1 General conditions for continual dynamics

We now prove the existence of continual evolution for the generic model represented by Eqs.(1). The main idea of the proof holds independently from context, and works in arbitrary trait spaces, both continuous and discrete.

We assume the existence of one-dimensional projections *u* ↦ *M* (*u*) and *φ* ↦ Φ(*φ*), both taking real values, and focus on studying the evolutionary dynamics in the (*M*, Φ)-plane. In general, it will not be possible to determine the values of *∂M/∂t* and *∂*Φ*/∂t* in terms of *M* and Φ, i.e. without knowing *u* and *φ*. However, to prove continual evolution one merely needs to control the *sign* of the two derivatives in suitable regions in the (*M*, Φ)-plane.

In the most general scenario, there is no specification on the projections *M* (*u*) and Φ(*φ*), nor is there any information on the dynamics in (*u, φ*)-coordinates. The assumptions refer directly to the signs of *∂M/∂t* and *∂*Φ*/∂t* in different parts of the dynamical plane. In this scenario, there is no reference to how these conditions can be obtained from knowledge of the dynamics in the (*u, φ*)-coordinates; the conditions are taken as given.

A fast positive feedback is modeled directly, by assuming that a nearly maximal value of *M* implies that *∂M/∂t* is positive, and similarly that a nearly minimal value of *M* implies that *∂M/∂t* is negative. We also assume the existence of a slowly reacting, negative feedback, modeled indirectly via Φ: for large *M* the external factor Φ will slowly increase, and for sufficiently large Φ the value of *M* will slowly decrease. Similar assumptions are stated for small *M* and small Φ. While the value of Φ adapts to the value of *M* very slowly, we assume that the negative feedback eventually dominates the positive feedback: for Φ sufficiently large the rate *∂M/∂t* is negative, no matter the value of *M*.

These feedback assumptions are stated more formally in SI section S1.2, conditions (PF1, PF2, NF1, NF2). The following result, which guarantees continual evolution, is proved there under these feedback assumptions, and provided there is a sufficiently strong separation of time scales between the ecological dynamics in *M* on the one hand, and the ecological dynamics in Φ and the effects of mutations on the other.

#### Theorem.

*If the feedback assumptions above are satisfied, then there exist initial values* (*u, φ*) *for which M* (*t*)*will fluctuate infinitely often between values arbitrarily close to both a maximum and a minimum value*.

Figure 1 illustrates the emerging behavior in the (*M*, Φ)-plane. When *M* is sufficiently close to (but, due to mutations, never equal to) its maximal value, Φ increases until it gets very close to a sufficiently large value Φ_+_, which in turn guarantees a decrease in *M* due to the dominant negative feedback. This decrease is initially small, while Φ is still increasing. After a limited amount of time, *M* eventually reaches a value arbitrarily close to its minimum and Φ decreases although, due to the timescale separation assumption, the decrease in Φ is arbitrarily small in this time interval, and hence the decrease in *M* is not prevented by Φ. The symmetry in the feedback conditions guarantees that the process repeats itself.

**Figure 1:**
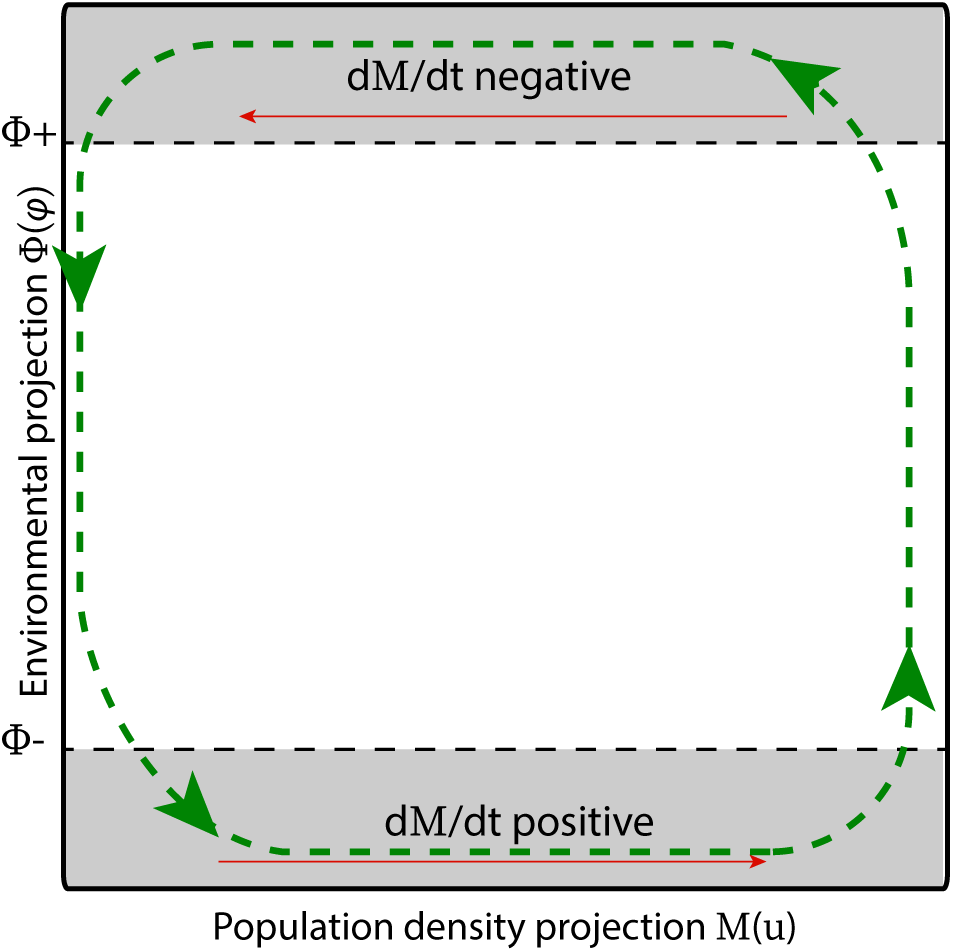
Sketch of emerging behavior in a 1-dimensional projection. The dynamics in the (*M* (*u*), Φ(*φ*)) plane, where *M* (*u*) is a one-dimensional projection from the population density space and Φ(*φ*) a one-dimensional projection from the environmental space. Because of the assumptions on the sign of the derivatives of these projections (red arrows), there will be continual dynamics (green arrows).

The simple formulation of the negative feedback conditions (NF1 and NF2) is not realistic on the whole (*u, φ*)-space. For example, when the population density *u* is concentrated around a single trait, one cannot expect any dynamics due to ecology, regardless of Φ, because near such single trait the rate of change *∂M/∂t* is governed almost entirely by mutations, which by assumption acts at a much slower time scale. Also, in the proof sketched above it is essential that the time interval needed for *M* to decrease to a value close to its minimum is independent of the slower time scale. It is sufficient, however, if the feedback conditions hold in the region of (*u, φ*)-space reached by the dynamics. In the more explicit case discussed below, we exploit this observation to obtain more realistic feedback assumptions.

### 3.2 Continual dynamics with a single evolving trait

Let us discuss a more explicit case involving a continuous 1-dimensional trait space, *x* (representing, for example, a continuous trait such as length) and a single environmental factor, *φ*. For this case, we will use as the one-dimensional projection *M* (*u*) the *average trait value*:

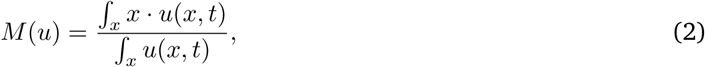

As explained in SI section S1.3, suitable feedback conditions in the (*u, φ*) space determine to a large extent the dynamical behavior in the (*M, φ*)-plane. While it will generally not be possible to deduce *∂M/∂t* and *∂φ/∂t* from only knowing *M* and *φ*, it will be possible to determine the signs of these rates in large regions of the plane. As discussed in the previous section, this can be sufficient for deducing continual evolution.

As a practical example, let us consider the logistic growth model, that is, that the growth function in Eqs.(1) is given by:

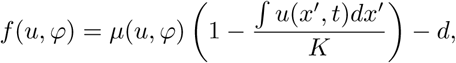

where *d* represents a mortality rate, *K* the carrying capacity, and the growth rate *µ* depends on the populations density *u* and environmental factor *φ*. The positive feedback is incorporated directly through the effect of *u* on the growth rate *µ*, while the negative feedback is incorporated indirectly through the effect of *u* on *φ* and the (opposite) effect of *φ* on *µ*. Figure 2 illustrates the indirect negative feedback.

**Figure 2:**
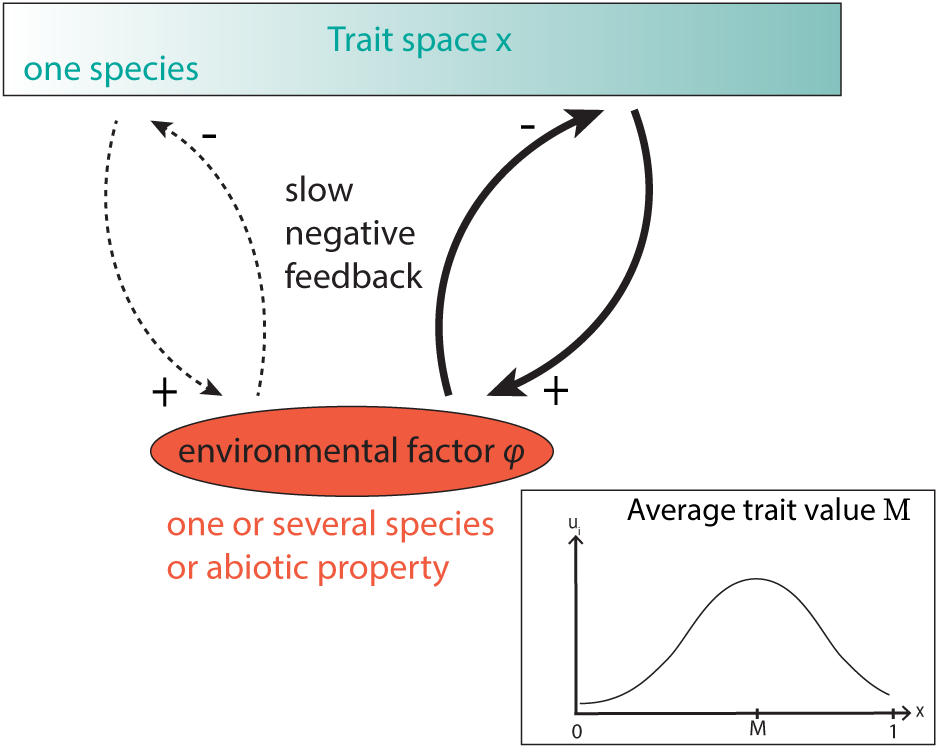
A single evolving trait with a range of possible trait values. One species has a single evolving trait, with a phenotype ranging from *x* = 0 to *x* = 1. Higher values of *x* denote a stronger interaction with the environment *φ* (which can be a species, an abiotic factor, or a property of the ecosystem). The interaction with the environment leads to the slow negative feedback. The inset illustrates that *M* is calculated by taking the average trait value in the population, thus weighing the trait values by the density of that phenotype (Eq. (2)).

As we show in SI section S1.3, given the mean trait value *M* (see Eq.(2)) the sign of 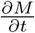 can be determined from the values of *M* and *φ* when *µ* is assumed to be either constant or strictly monotonic for all relevant parameters. Therefore, under the corresponding feedback assumptions and time scale separation assumptions, the logistic growth model can satisfy continual evolutionary dynamics.

To simulate this example numerically, we split the population in discrete groups with similar trait values. Figure 3A shows that forcing the absence of evolution leads to extinction of all but one groups; fast evolution (i.e. large *ϵ*_*e*_) leads to an equilibrium, that is, to a static *u*(*x, t*) that shows one single, well-defined mean value *M*; and slow evolution leads to continual evolutionary fluctuations where the average trait value alternatingly approaches one of the two extremal values. Figure 3B shows the distribution of phenotypes over time for the latter case. The population is centered around one of the extreme trait values most of the time, but intermediate phenotypes are also seen in the transition periods. Figure 3C shows that the diversity of phenotypes, measured by the Shannon index (SI section S2.1), peaks at shifts of the mean trait value due to the flattening of *u* during the transition period, which increases the standard deviation of the distribution (see Bonachela et al. ^17^).

**Figure 3:**
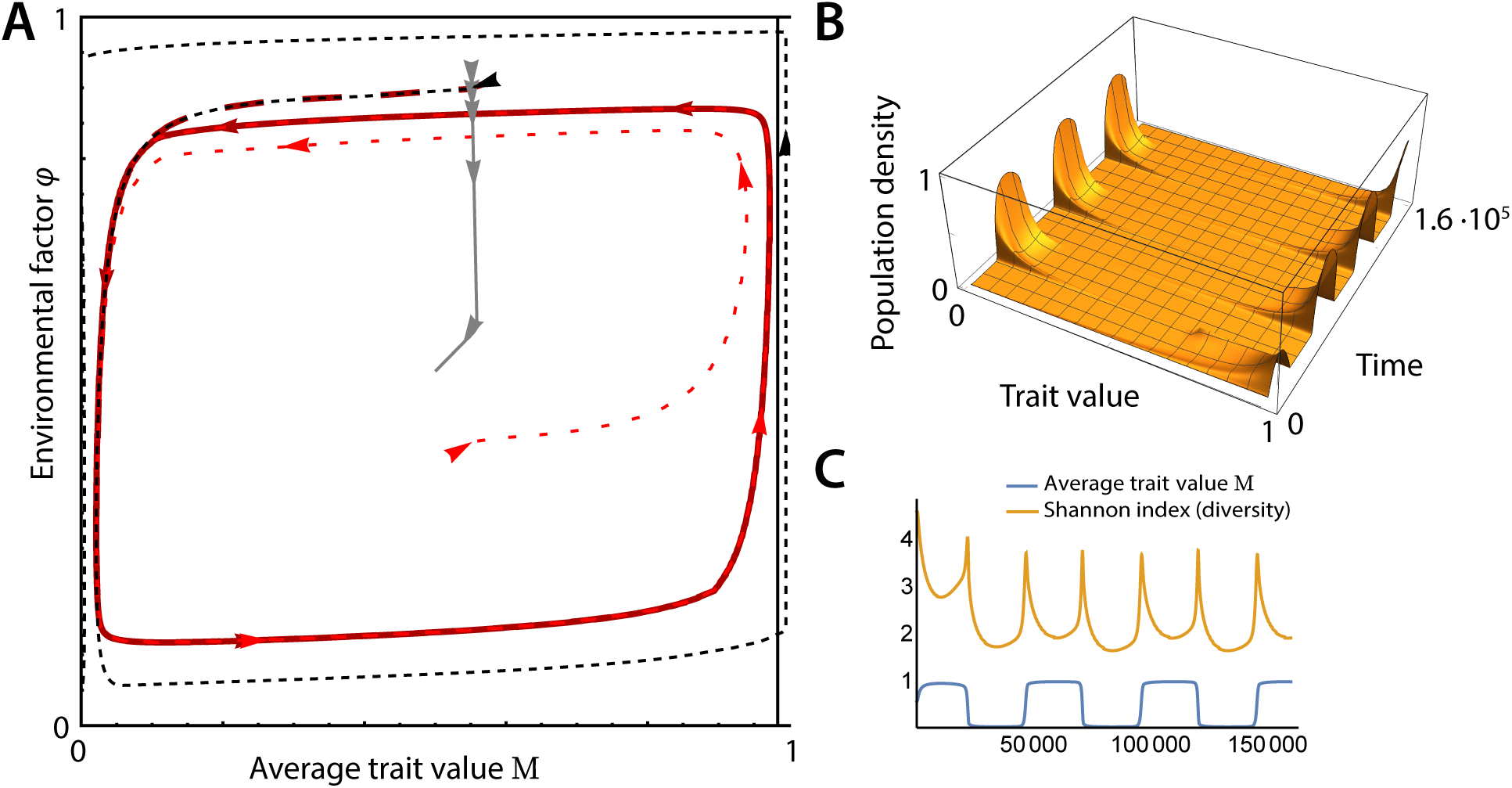
Example of evolutionary dynamics with a single evolving trait. Behaviour of a system following the outline of Figure 1. **A** System behaviour for no evolution (thin dashed lines, resulting in the extinction of the population), fast evolution (gray dashed line, leading to an equilibrium), and slow evolution (red and solid dashed lines; continual evolutionary dynamics from different initial values). **B** Phenotype abundances obtained in the continual evolution case. **C** Mean trait value and Shannon diversity index for the same case, showing the emerging fluctuations. Simulation of 100 phenotypes equally distributed over the range of trait values; see SI section S2.3 for the exact equations and parameter values.

### 3.3 Conditions for continual dynamics in the case of a single evolving trait consisting of two phenotypes

To understand in more detail the role of a fast positive and slow negative feedback in the emergence of the evolutionary oscillations, we reduced the system to a simpler version. In this case, the trait space *x* consists of only two points, *A* and *B*, representing a trait with only two main values (i.e. only two phenotypes are possible). One of the phenotypes is interacting with a external factor and the other one either interacts less strongly or not at all (see Figure 4A). This representation reflects a biological system where a trait value is either present or not at all, e.g. a parasite choosing one host type over another. When the trait also affects mate choice (e.g. because they will be located near the same host), the trait has a positive feedback. The slow negative feedback could result from the fact that one of the hosts can develop defenses.

**Figure 4:**
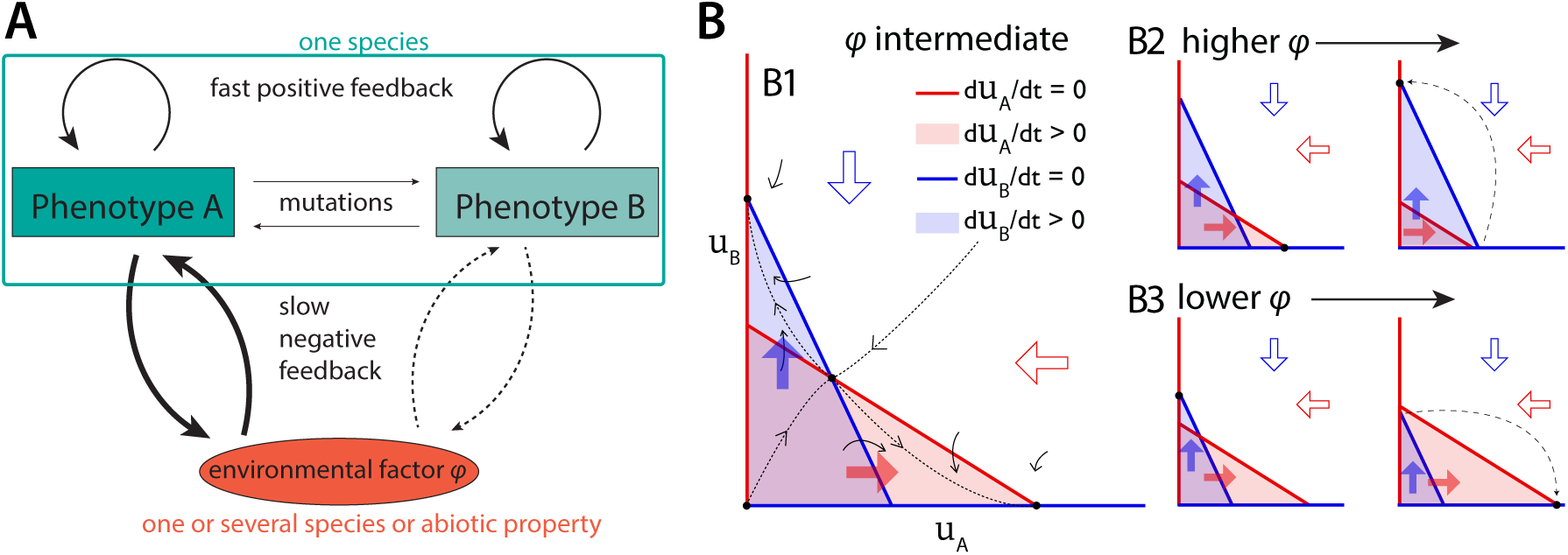
Continual evolution with a single evolving trait consisting of two phenotypes. **A** A species with only two possible phenotypes, *A* and *B*, which both show fast positive feedbacks on their own phenotype. Phenotype *A* shows a strong negative feedback on an external factor *φ*, while phenotype *B* shows a weaker, negative feedback (or none at all). Mutations are only possible between the two phenotypes, although rates are low. **B** Approximate phase planes. Block arrows and colored areas show the sign of the derivatives. Blue and red lines show the isoclines where the derivatives are 0. The isoclines for *du*_*A*_*/dt* = 0 and *du*_*B*_*/dt* = 0 cross as shown, because the fast positive feedback ensures that the growth of the subpopulation *u*_*B*_ decreases faster with increasing *u*_*A*_ than the growth of the subpopulation *u*_*A*_, and vice versa. Black dashed lines show the direction of change over time. Different phase planes with black solid lines above show how the phase planes will change over time.

We analysed this system, where individuals with both phenotypes (*u*_*A*_ and *u*_*B*_) and the environmental factors (*φ*) change over time. Phenotype *A* could represent e.g. the parasite phenotype choosing the host that can develop defenses; the (less) interacting phenotype *B* could represent the phenotype choosing the host that cannot develop defenses, and the external factors *φ* would therefore refer to the amount of defense (for the host that can develop defenses). Thus, *φ* is positively correlated with *u*_*A*_ and negatively correlated with *u*_*B*_:

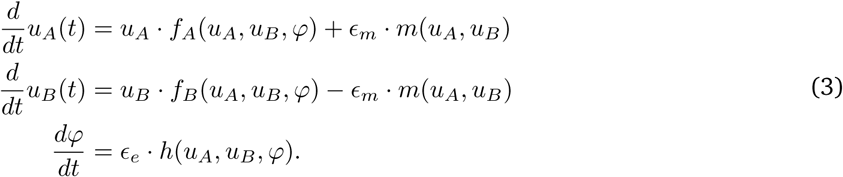

This set of equations directly results from Eqs(1) by splitting the population density distribution in two subpopulations (individuals showing trait value *A*, i.e. phenotype *A*, and those with phenotype *B*). Because the only relevant mutations in this case are mutations that change the phenotype from *A* to *B* and vice versa, the evolution function *g* can be simplified into a function *m* that describes the net mutations from one trait value to the other (hence the opposite sign). The dependency of the functions *f*_*A*_ and *f*_*B*_ on the subpopulations *u*_*A*_ and *u*_*B*_ is assumed to cause a positive feedback (see condition (PF) in SI section S1.1), while the dependency on *φ* of the functions *f*_*A*_, *f*_*B*_, and *h* generate a negative feedback, given by conditions (NF1) and (NF2), assumed to eventually dominate over the positive feedback. In SI section S1.1, we prove that, under the additional, mild “unique stable value assumptions” (USV1 and USV2) a sufficiently small *ϵ*_*m*_ and *ϵ*_*e*_ lead to the emergence of continual evolution. It is both natural and necessary to assume that the effect of evolution is relatively small (but never zero), as large values of *ϵ*_*m*_ typically cause convergence to an equilibrium. The detailed proof can be found in SI section S1.1.

The idea of the proof for a variable population size is illustrated in Figure 4B. The diagrams show an approximation of the dynamics in the (*u*_*A*_,*u*_*B*_) phase planes. This approximation is without the mutation term, but since mutations are rare (in our case the mutation term is small), the actual phase plane will not be very different. The assumptions above guarantee that the phase plane for intermediate *φ* resemble the one shown in panel B1. When the environmental strength *φ* is intermediate, there are four intersections of the lines 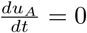 and 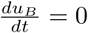, i.e. four equilibria, one of which is repelling (the origin), another one is a saddle point (the point in the middle), and the other two are attractors. The nature of the equilibria can be seen from the sign of the derivatives. Depending on the initial value for *u*_*A*_ and *u*_*B*_, the system converges to an attractor near the *u*_*A*_-axis or the *u*_*B*_-axis. When the system is near the former, *u*_*B*_ will be almost 0 and since *u*_*A*_ has a positive effect on *φ, φ* will increase. The change in *φ* will in turn change the phase plane diagram and the intersection near the *u*_*A*_-axis will become repelling (panel B2). When *φ* is high enough,there will be only one attracting intersection (near the *u*_*B*_-axis), and the system will approach that state. As *u*_*B*_ increases (and *u*_*A*_ decreases) *φ* decreases, but, since the fast-slow condition guarantees that the dynamics in *φ* are slower than in *u*_*A*_ and *u*_*B*_, the system will come close to the intersection near the *u*_*B*_-axis before *φ* changes significantly. When *φ* does change, the system will pass by panel B1 (figure for intermediate *φ*), but now *φ* will continue to decrease and the intersection near the *u*_*B*_-axis will eventually disappear (panel B3). This cycle will continue indefinitely, giving rise to continual evolution.

The proof of continual dynamics above only required some restrictions on the functions *f*, *m* and *h* and did not use any functional forms. However, to give an indication of the expected dynamics, we constructed an example that follows these restrictions (see Figure 5 and SI section S2.2 for details and an additional example). Our proof states that stable continual evolutionary cycles will emerge for almost all initial values if *ϵ*_*e*_ and *ϵ*_*m*_ are chosen to be sufficiently small. Figure 5 shows that this is the case by using two different initial states. On the other hand, a high mutation rate and/or a fast feedback lead to a stable equilibrium (i.e. stasis), whereas the absence of mutations causes one of the two phenotypes to go extinct.

**Figure 5:**
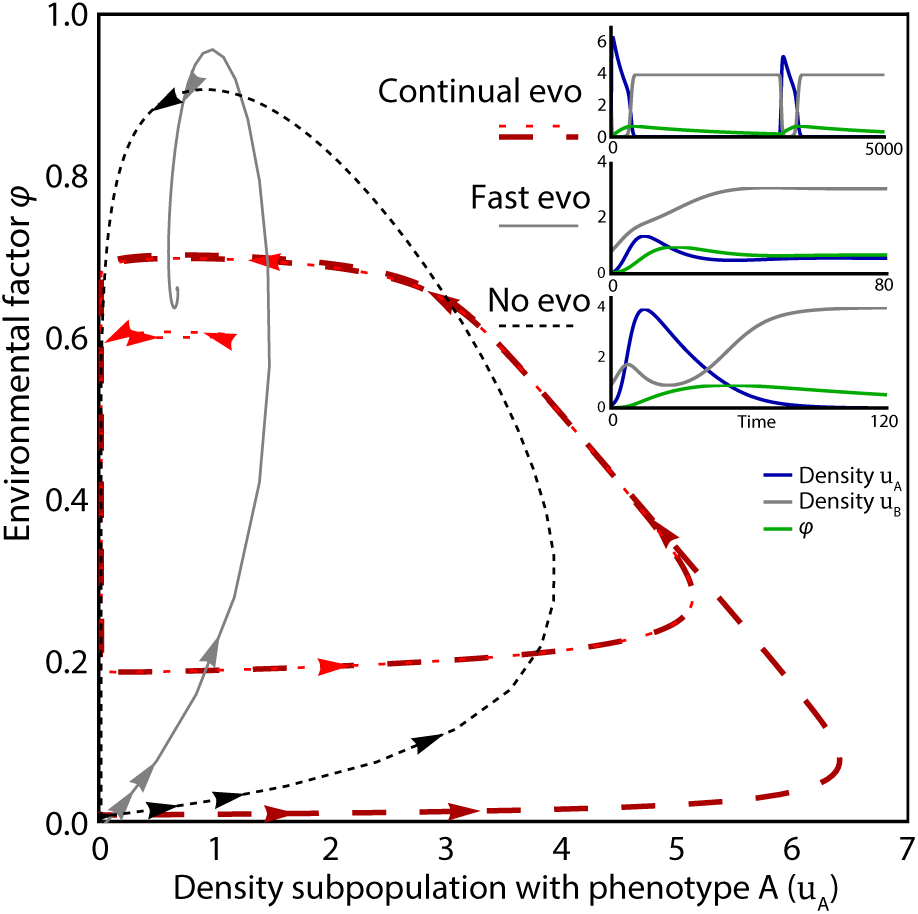
Example of evolutionary dynamics with a single evolving trait consisting of two phenotypes. Behaviour of a specific system following the outline of Fig 4A. Phase plane diagram of the density of individuals with phenotype A and environment *φ* shows the possible system behaviour for different values for the relative timescales of population dynamics, feedbacks, and mutation. Slow feedback and mutations (*ϵ*_*m*_ = 0.00005 and *ϵ*_*e*_ = 0.0005) lead to continual cyclic evolutionary dynamics (red lines and top inset), almost independent of initial conditions (two initial conditions shown). With fast feedback and mutations (*ϵ*_*m*_ = 0.01 and *ϵ*_*e*_ = 0.1), an equilibrium is reached (gray line and middle inset). No evolution (*ϵ*_*m*_ = 0 and *ϵ*_*e*_ = 0.01) leads to the extinction of one of the traits (black dashed line and bottom inset). See SI section S2.2 for equations and parameter values.

### 3.4 Continual dynamics in the case of two evolving traits: an example

In natural systems, multiple traits determine the fitness of an individual. An extension of our model to more traits can thus provide a more realistic picture of the dynamics expected in natural populations. We extended the system to a species with two evolving traits, each associated with a different environmental factor through a negative feedback. In other words, the trait space *x* is now a two-dimensional space and, since both traits are associated with a different environmental factor, the space of external factors *y* is also a two-dimensional space.

For the sake of simplicity, we discretised the trait space, and therefore the number of possible phenotypes is now the product of the number of phenotypes for each of the traits. In the simplest case, which we only use for a schematic representation of the population, there are only two values per trait, thus defining four possible phenotypes in the population (Figure 6A): high susceptibility for both feedbacks (A1 in that figure), low susceptibility for both feedbacks (B2), and high susceptibility for one and low for the other feedback (A2 and B1). For our simulations, we extended the system to eight possible values per trait, therefore 64 possible phenotypes in total. Our results show that this system can lead to continual evolution. Interestingly, the emerging evolutionary fluctuations generally become irregular (in Fig. 6B, shown through the mean trait value *M* for each trait of the population). Moreover, looking at the change of the diversity index over time, we can conclude that there is no fixed period in the dynamics (Figure 6C). The example with two traits thus shows that we do not expect a natural system to come back to exactly the same state recurrently. See SI section S2.4 for more details.

**Figure 6:**
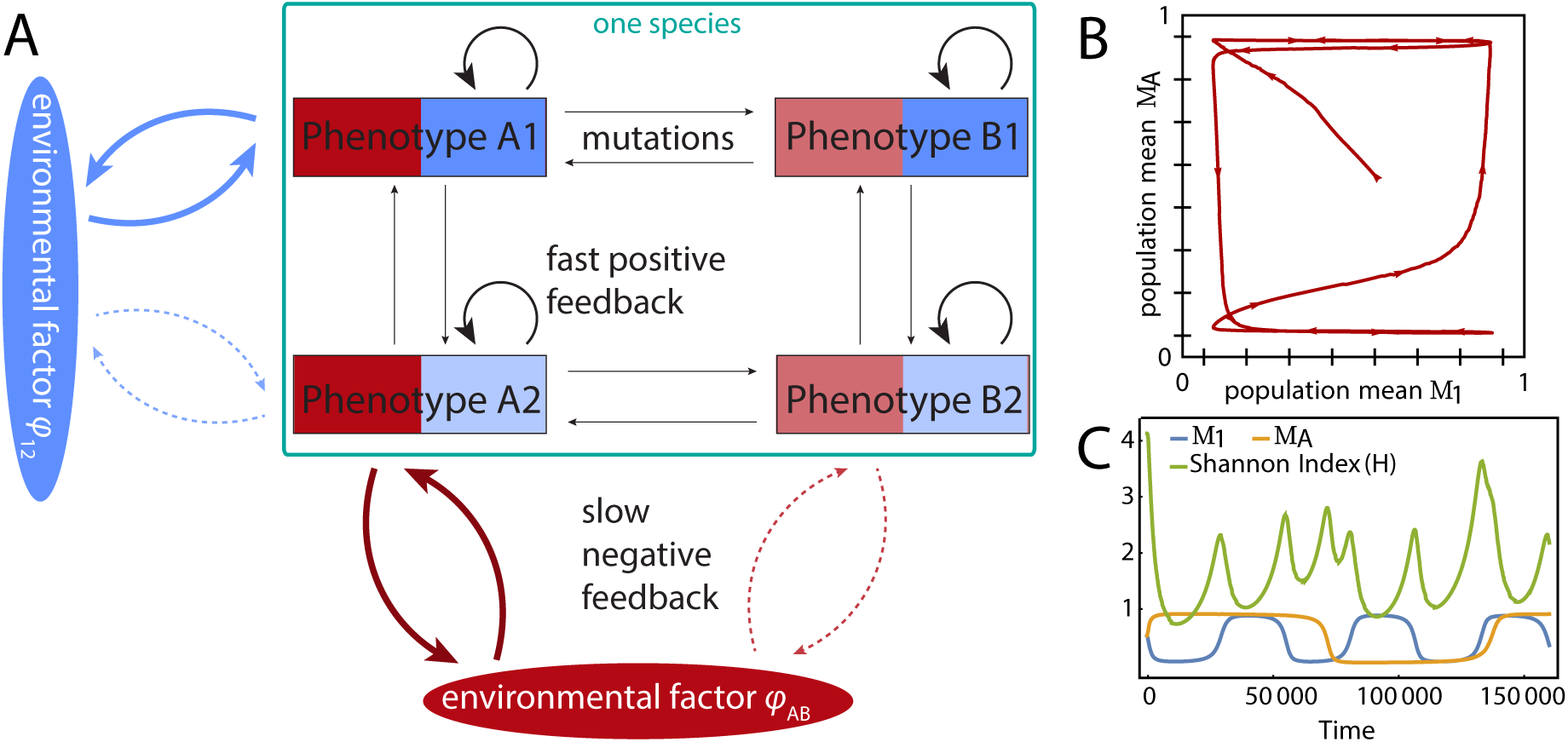
Example of evolutionary dynamics with two evolving traits. **A** We consider two environmental factors (*φ*_12_ and *φ*_*AB*_, where the character of the subscript refer, respectively, to the phenotype the factor affects strongly and weakly) and, in the simplest case, each trait consists of two trait values, leading to four phenotypes in total. More phenotypes per trait are also possible, leading to a total number of phenotypes as the product of the possible values per trait. **B** The change in the subpopulation-level mean value for each trait (*M*_1_ for one trait and *M*_*A*_ for the other trait) in time for an example with eight possible phenotypes per trait (indicated by the tics on the axes) and therefore 64 phenotypes in total. **C** Mean trait values and diversity over time. With more than one trait, periods are irregular as can be seen from the non-periodic Shannon Index (*H*). Equations and parameters for the figure are given in the SI section S2.4.

## 4 Discussion

We have shown that, when allowing for phenotypic variation in the population, a simple motif of fast positive and slow (but eventually dominant) negative feedbacks leads to continual evolution regardless of whether mutations have small or large effects. Our results are general because the evolutionary dynamics do not depend on specific equations, parameter values, or dependencies between traits (trade-offs). Our results are also robust to small temporal change in the population density distribution (e.g. migration, random effects). Previous work has shown that continual evolution can be found in specific (and typically simple) models. In the literature, generality is typically claimed by showing that the reported results remain for a range of parameter values or different equations ^20,11^. Here, we avoided the use of specific equations and parameter values to obtain truly general results that are robust across parameter values and functional forms. On the other hand, previous examples that have avoided using specific functional forms to explain some aspect of continual evolution required additional restrictions on the models. For example, Dercole et al. ^20^ focussed on predator-prey interactions, Nordbotten and Stenseth ^9^ considered bilinear species interactions, and Bonachela et al. ^17^ considered a fixed set of three species that interact forming a non-transitive cycle. Our proofs are accompanied by specific examples for the sake of concreteness, but our results are not restricted to those examples.

In all our examples, continual evolutionary dynamics stem from switching between multiple ecological attractors that are nodes (ecogenetically driven RQ dynamics ^21^). Our conclusions remain the same when these attractors are limit cycles instead. An example of this type of dynamics is given in Khibnik and Kondrashov ^21^ Fig. 4. Interestingly, including a polymorphic population in their example removes the evolutionary oscillations, i.e. only ecological cycles remain, which could be equivalent to ecologically driven RQ dynamics, that is, fluctuations in traits that are fast and small and only follow ecological population density oscillations. However, when we make their negative feedback slower by decreasing the parameters for the predator dynamics tenfold (their parameters r4 and *γ*), ecogenetically driven RQ dynamics are retrieved (results not shown), which supports the applicability of our claims to existing examples.

Some of our results are in line with and extend previous conclusions. The dynamics of continual evolution are very similar to microevolutionary RQ dynamics; the difference is that continual evolution does not require co-evolution. Khibnik and Kondrashov ^21^ mention that RQ dynamics with a single evolving trait are possible when the dynamics are ecologically driven (the traits follow ecological dynamics), or ecogenetically driven and switching between two different ecological attractors. Continual evolution of a single trait might be quite prevalent, and examples include predator-prey systems in which the prey evolves but the predator does not (or much more slowly, as might be relatively common, see e.g. Vermeij ^22^). Moreover, our case of a single evolving trait with two phenotypes reproduces previous results by Mougi and Iwasa ^11^ showing that fast adaptation is less likely to lead to RQ dynamics. In Bonachela et al. ^17^, we also found that biotic drivers that respond too fast to the environment do not lead to RQ dynamics and, similar to that paper the dynamics in this work are not linked to population density fluctuations. Also, as observed before (e.g. Bonachela et al. ^17^), we show for a 1-dimensional trait space that evolution should not be too slow relative to the negative feedback, and that ecology and evolutionary dynamics can influence each other to guarantee continual dynamics. The average population trait has to change drastically while the negative feedback changes relatively little. If a trait evolved gradually, evolution should not be much slower than the dynamics of the feedback or otherwise the feedback would ‘catch up’ with the average population trait while this average is near the equilibrium value. An example of such kind of dynamics could be found in predator-prey systems with a slowly evolving predator (due to, e.g. generation times that are much longer for the predator than for the prey, like in the laboratory system of an algae and a rotifer ^23^). In these cases, non-optimal predator phenotypes might stay in the population and not go extinct. Thus, the species does not need de novo mutations for continual evolution to emerge, as the trait space can be explored through selection and recombination. This type of RQ dynamics where the traits stay in the population are more common in higher organisms, and is an example of evolutionary rescue. Faster evolutionary change through mutations might also occur when species are regularly exposed to different environments and evolve adaptability ^24,25^.

We included the possibility of a polymorphic population, often left out of RQ dynamics studies and which can have a pronounced effect on the results. As shown in SI section S2.5, examples existing in the literature of models showing RQ dynamics can lose these evolutionary oscillations when the more general setting of a polymorphic population is considered. Even when the original models do not seem to be directly linked to the motif described in this paper, including our fast positive-slow negative feedback combination leads to recovering continual evolution. This the case for models such as the one in Mougi and Iwasa ^11^, where the RQ dynamics remain when polymorphism is introduced (data not shown) but are now ecologically driven (i.e. evolution just follows the population’s ecological predator-prey cycles). Other cases that do include polymorphic populations are the stochastic model by Dieckmann et al. ^10^; models of evolutionary branching and extinction using adaptive dynamics (summarized in Kisdi et al. ^16^); and a Lotka-Volterra predator-prey model with a polymorphic population that, similarly to our results, shows examples of RQ dynamics ^26^.

On the one hand, collecting evidence of continual evolution from natural systems is a difficult task ^18^ and, in experiments, the need for long term measurements is a problem. One experiment that shows the effect of evolution on predator-prey dynamics suggests RQ dynamics ^11^ but, due to the limited measurement time, it cannot be excluded that the cycles will eventually dampen. On the other hand, there is an increase in long term adaptation experiments, but these are usually in bacterial systems whereas most of the modelling has focused on sexually reproducing predator-prey communities. Here, microbial systems are included by allowing for polymorphism and larger mutation effect sizes. In bacterial systems polymorphic populations are common even under reasonably constant conditions as recently shown in the Long Term Evolution Experiment with *E. coli* ^27^. A polymorphic trait distribution might result from density dependent dynamics, as shown theoretically in a chemostat ^28^. Therefore, our results are an important addition for linking theoretical to experimental observations of evolutionary dynamics.

Although we have tried to keep our framework general, our results have some limitations. We show that, under specific assumptions, we can guarantee the emergence of continual evolution. Systems that do not follow these assumptions (e.g. fast negative feedbacks, no positive feedback, not enough timescale separation) are more likely to lead to stasis. However, we did not prove any conditions that necessarily lead to stasis. Also, we did not include individual variation within the population. Future work will focus on combining the methods used in this paper with a polymorphic population to link the ecological and evolutionary timescales with individual-based models. In addition, our results are limited to only one evolving species, although the feedbacks we describe could come from another evolving species or be a result of a more complex ecosystem. A next step would thus be extending our study to more than one evolving species. Finally, the results in this paper are constrained to micro-evolutionary dynamics, but the general model (Eq. (1)) allows for extension to macro-evolutionary phenomena. In this respect, our results are in line with the results in Doebeli and Ispolatov ^29^, which show that most coevolutionary dynamics are found with intermediate diversity, where not all niches are filled. Here we see that, if diversification emerges (and, therefore, a phenotypically polymorphic population that fills more niches), the co-evolutionary dynamics cease (see SI Figure S3).

In conclusion, our general framework shows, without constraining functional forms nor phenotypic diversity within the system, that a fast positive feedback combined with a slow negative feedback leads to continual dynamics if there is enough separation between the timescales involved. By doing so, we have improved the understanding of continual evolution and co-evolution in a large class of systems. Importantly, our general framework may be used to study and predict evolutionary dynamics without knowledge of the exact equations describing the system, and thus assess what likely happens and has happened in nature.

## 5 Acknowledgements

The authors thank Jan Nordbottom and Øistein Haugsten Holen for helpful discussions and comments on the manuscript. MTW and NCS were supported by 227860/F20 ERC-Stenseth “Red Queen coevolution in multispecies communities: long-term evolutionary consequences of biotic and abiotic interactions” and CEES-core funding Research Council of Norway through the Centre for Ecology and Evolutionary Synthesis.

## 6 Author contributions

MTW, HP and NCS designed the research. MTW developed the models. HP developed the mathematical proofs. MTW and JAB wrote the manuscript with input from HP and NCS.

