## Supplementary Information for "Continual evolution through coupled fast and slow feedbacks"

### Contents

S1. Rigorous Formulation and Proofs 1
S1.1. One evolving trait with two phenotypes 1
S1.2. General case: Projection to two dimensions 5
S1.3. One evolving trait with a continuous range of phenotypes 7
S2. Examples: Rate equations, diversity index and supplementary figures 10
S2.1. Shannon index 10
S2.2. One evolving trait with two phenotypes 10
S2.3. One evolving trait with a range of phenotypes 12
S2.4. Two evolving traits 12
S2.5. Literature models 13
S3. Supplementary Tables 13
References 14

### S1. Rigorous Formulation and Proofs

We first treat the simplest case of one evolving trait with only two different phenotypes (Section S1.1). We then show how to adapt the key ideas to obtain a general statement, valid for an arbitrary trait space $x$ and environmental space $y$ (Section S1.2). In Section S1.3 we treat a more explicit example, given by the logistic model involving one trait with a continuous range of phenotypes, and show that the ideas of the general case can be adapted to this more explicit setting.

#### S1.1. **One evolving trait with two phenotypes.**

We first consider the simplest possible system where the combination of a slow negative and a fast but weaker positive feedback cause continual evolution. We consider a single species with only two phenotypes $A$ and $B$ (we refer to the population densities of the subpopulation with phenotype $A$ and $B$ with just $A$ and $B$, short for $u(A)$ and $u(B)$), and we will introduce a negative feedback induced by a single environmental factor $\varphi$. We will write

$$R = \frac{A}{A+B}.$$

Thus $R$ takes on values in the interval $[0, 1]$. We assume that the total population will remain bounded, from above as well as below, so that $R \to 0$ corresponds to the extinction of population $A$, and $R \to 1$ to the extinction of population $B$. In describing the differential equations governing the dynamics of $A$, $B$ and $\varphi$, we separate the ecological factors and the evolutionary factors. The rate of change in $A$ due to ecological factors naturally vanishes when $A = 0$, and similarly the rate of change of $B$ due to ecological factors vanishes when $B = 0$. Thus we can write the system of differential equations in the form

$$\frac{dA}{dt} = A \cdot f_A(A, B, \varphi) + \epsilon_m \cdot g(A, B),$$
$$\frac{dB}{dt} = B \cdot f_B(A, B, \varphi) - \epsilon_m \cdot g(A, B), \quad \text{and}$$
$$\frac{d\varphi}{dt} = \epsilon_e \cdot h(A, B, \varphi),$$

where $f_A$ and $f_b$ represent ecological factors and $g$ represents the rate of change due to mutations.

We will assume that the functions $f_A, f_B, g$, and $h$ are all continuously differentiable, and the constants $\epsilon_m$ and $\epsilon_e$ will be chosen arbitrarily small in order to represent that both mutations and the change in the environmental factor $\varphi$ occur at a slower time scale than the ecology of $A$ and $B$. The function $g$ is assumed to be strictly positive for $A$ small, and strictly negative for $B$ small. Recall that the total population size is assumed to stay bounded from below, hence when $A$ is small $B$ is not, and vice versa.

The following assumption guarantees a positive ecological feedback on the subpopulations $A$ and $B$:

(PF) For all values of $A, B$, and $\varphi$ we have

$$\left(\frac{\partial}{\partial A} - \frac{\partial}{\partial B}\right) f_A > 0, \quad \text{and} \quad \left(\frac{\partial}{\partial B} - \frac{\partial}{\partial A}\right) f_B > 0.$$

Thus, if an amount of $B$ is replaced by an equal amount of $A$ while $\varphi$ remains constant, the fitness of the population $A$ increases while the fitness of the population $B$ decreases. It is clear that in absence of mutations and in an environment with a constant environmental factor $\varphi$ the (PF) assumption would lead to extinction of either $A$ or $B$ for almost all initial values.

The next assumption, which we will refer to as the *unique stable value assumption*, will be used to draw conclusions about the sign of $\frac{dR}{dt}$ without knowing the exact values of $A$ and $B$. We do not claim that this assumption is necessary for our results, but it turns out to be satisfied in many models and is certainly convenient:

(USV1) For fixed values of $B$ and $\varphi$, there is a unique value $A_0 \geq 0$ such that

$$f_A(A, B, \varphi) > 0$$

for $0 \leq A < A_0$, and

$$f_A(A, B, \varphi) < 0$$

for $A > A_0$.

(USV2) For fixed values of $A$ and $\varphi$, there is a unique value $B_0 \geq 0$ such that

$$f_B(A, B, \varphi) > 0$$

for $B < B_0$, and

$$f_B(A, B, \varphi) < 0$$

for $B > B_0$.

For $\varphi$ fixed we can therefore consider $A_0$ as a graph $\eta_A(B, \varphi)$, and similarly $B_0 = \eta_B(A, \varphi)$. The positive feedback condition guarantees the following:

**Lemma 1.** *The derivatives* $\frac{\partial \eta_A}{\partial B}$ *and* $\frac{\partial \eta_B}{\partial A}$ *are both strictly less than* $-1$*, and possibly* $-\infty$.

*Proof.* We prove the first statement, the second is analogous. Let $A_0, B, \varphi$ be a triple for which $f_A(A_0, B, \varphi) = 0$. The unique stable value assumption implies that

$$\frac{\partial f_A}{\partial A}(A_0, B, \varphi) \leq 0,$$

and the positive feedback assumption gives

$$\frac{\partial f_A}{\partial B} < \frac{\partial f_A}{\partial A}.$$

The function $\eta_A$ is implicitly defined by $f_A(\eta_A(B, \varphi), B, \varphi) = 0$. When $\frac{\partial f_A}{\partial A} \neq 0$ the Implicit Function Theorem gives

$$\frac{\partial \eta_A}{\partial B} = -\frac{\frac{\partial f_A}{\partial B}}{\frac{\partial f_A}{\partial A}}$$

which gives $\frac{\partial \eta_A}{\partial B} < -1$. When $\frac{\partial f_A}{\partial A} = 0$ we have $\frac{\partial f_A}{\partial B} < 0$ by assumption (PF), and hence $\frac{\partial \eta_A}{\partial B} = -\infty$. □

We now will impose three conditions that guarantee a negative feedback induced by the environmental factor $\varphi$. The first of these conditions is clear: replacing $B$ with $A$ is beneficial for $\varphi$, while an increase in $\varphi$ is beneficial for $B$. The second condition guarantees that the negative feedback dominates the positive feedback for extreme values of $\varphi$.

(NF1)
$$\frac{\partial}{\partial \varphi}(f_A - f_B) < 0 \quad \text{and} \quad \left(\frac{\partial}{\partial A} - \frac{\partial}{\partial B}\right) h > 0.$$

(NF2) There exists $0 < \varphi_{--} < \varphi_{++} < 1$ such that

$$f_B > f_A$$

whenever $\varphi > \varphi_{++}$, while

$$f_A > f_B$$

whenever $\varphi < \varphi_{--}$.

Hence, for $\varphi$ sufficiently large and $\varphi$ sufficiently small the level sets $\{f_A = 0\}$ and $\{f_B = 0\}$ do not intersect. We may write $\varphi_{++}$ and $\varphi_{--}$ for the maximal resp. minimal value of $\varphi$ for which the level sets intersect.

Observe that condition (NF2) would be vacuous if the values $\varphi_{--}$ and $\varphi_{++}$ could never be reached. In order to guarantee that the negative feedback eventually dominates the positive feedback we therefore assume a final condition, which we will refer to as a "transitivity" assumption:

(Tr) The function $h$ is strictly positive when $\varphi \leq \varphi_{++}$, $B = 0$ and $A = \eta_A(0, \varphi_{++})$. Similarly, the function $h$ is strictly negative when $\varphi \geq \varphi_{--}$, $A = 0$ and $B = \eta_B(0, \varphi_{--})$.

Let us consider, for a fixed value of $\varphi$, joint solutions of the two equations

$$A \cdot f_A = 0 \text{ and } B \cdot f_B = 0.$$

It is clear that, besides the origin, there always is a unique solution on each of the axes. For $A, B > 0$ it follows from Lemma 1 that there either is no solution or a unique solution, depending on the value of $\varphi$. For $\varphi = \varphi_{++}$ the intersection point of $\{f_A = 0\}$ and $\{f_B = 0\}$ lies in the axis $\{A = 0\}$, while for $\varphi = \varphi_{--}$ the intersection point lies in the axis $\{B = 0\}$. See Figure S1.1 (left) for a simple depiction of the level sets $\{f_A = 0\}$ (in red) and $\{f_B = 0\}$ (in blue).

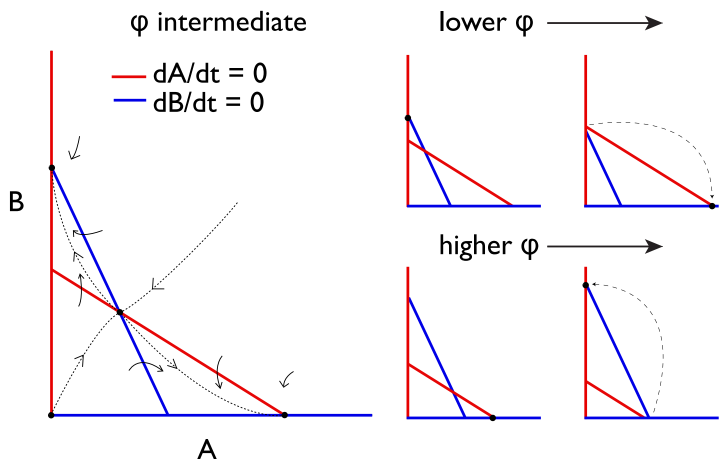


FIGURE S1. Left: Dynamics in the $(A, B)$-plane for fixed $\varphi_{--} < \varphi < \varphi_{++}$. Right: Changes in the phase plane when $\varphi$ decreases below $\varphi_{--}$ (top) or increases above $\varphi_{++}$ (top).

Let us consider the ecological dynamics in the $(A, B)$-plane caused by the differential equations

$$\frac{dA}{dt} = A \cdot f_A,$$
$$\frac{dB}{dt} = B \cdot f_B$$

for a fixed value of $\varphi$. When $\varphi \le \varphi_{--}$ or $\varphi \ge \varphi_{++}$ there are three fixed points where

$$\frac{dA}{dt} = \frac{dB}{dt} = 0,$$

namely the origin and the two intersection points of the curves $\{f_A = 0\}$ and $\{f_B = 0\}$ with the respective axes $\{B = 0\}$ and $\{A = 0\}$. The origin is always repelling. One of the points on the axes is a saddle fixed point, with stable manifold equal to the axis. The third fixed point is attracting, and all orbits of initial values not lying on the axes converge to this attracting fixed point.

When $\varphi_{--} < \varphi < \varphi_{++}$ there are four fixed points. Again the origin is a repelling fixed point. There are again two fixed points on the axes, which are now both attracting. Finally, there is an intersection point of the curves $\{f_A = 0\}$ and $\{f_B = 0\}$, and assumption (i) implies that this is a hyperbolic saddle fixed point. Its stable manifold is the separatrix of the two attracting basins.

Let us now consider the effect of the mutations, represented by $\epsilon_m \cdot g(A, B)$, on the dynamics in the $(A, B)$-plane in the case $\varphi_{--} < \varphi < \varphi_{++}$. The behavior near each of the fixed points is stable under small $C^1$-perturbations, and the qualitative behavior of the system is robust. Hence, by choosing $\epsilon_m$ sufficiently small, there will still be a repelling fixed point at the origin, and a saddle point with separatrix near the intersection point of the curves $\{f_A = 0\}$ and $\{f_B = 0\}$. The rest of the quadrant is attracted to neighborhoods of the original attracting fixed points. These neighborhoods can be chosen arbitrarily small by choosing $\epsilon_m$ sufficiently small. Note that the addition of mutations causes the axes to be repelling, hence the attracting fixed point no longer lies on the axis but sufficiently nearby.

Finally let us consider the full three-dimensional dynamical system, taking into account that $\varphi$ is not fixed. We assume that for given $t_0$ we have $\varphi(t_0) < \varphi_{++}$, and that $(A(t_0), B(t_0))$ lies in the small attracting region near the $A$-axis. For any $\delta > 0$ we can, by taking $\epsilon_e$ and $\epsilon_m$ sufficiently small, assume that $A(t_0) - \eta_A(0, \varphi_{++}) > -\delta$ and $B < \delta$. It follows that $h(A(t_0), B(t_0), \varphi(t_0))$ is strictly positive. It follows that $\varphi$ will continue to grow while $\varphi < \varphi_{++} + \delta$ and while $(A(t), B(t))$ remains trapped in the small attracting neighborhood near the $A$-axis.

Let us consider the first time $t_1$ at which either $\varphi \geq \varphi_{++} + \delta$ or at which $(A, B)$ leaves the small attracting neighborhood. In either case it follows that the orbit $(A(t), B(t))$ is guaranteed to approach the small attracting neighborhood near the $B$-axis. If $\epsilon_e$ is sufficiently small the value of $\varphi$ can only decrease arbitrarily little while this happens. The conclusion is that we end up with a time $t_2 > t_1$ when $\varphi(t_1) > \varphi_{--}$ and $(A(t_2), B(t_2))$ lies in the attracting neighborhood near the $B$-axis. By the symmetry of our assumptions the process will repeat itself. We have proved the following:

**Theorem 2.** *Let $0 < a < b < 1$, $\varphi_{--} < c < d < \varphi_{++}$, and write $K$ for the rectangle $[a, b] \times [c, d]$. Then for $\epsilon_e$ and $\epsilon_m$ sufficiently small there exists an orbit $(A(t), B(t), \varphi(t)$ for which the coordinates $(R(t), \varphi(t))$ avoid $K$, and for which $R(t)$ fluctuates between values larger than $b$ and smaller than $a$.*

S1.2. **General case: Projection to two dimensions.** Before considering systems with more (possibly infinitely many) traits, let us take look back at the strategy used in the previous section. We introduced a one-dimensional projection $(A, B) \mapsto R$, which happened to be the ratio between the subpopulation sizes $A$ and $B$. It turned out that in order to conclude continual evolution we needed to deduce the sign of $\partial R/\partial t$ in an appropriate region of $(A, B, \varphi)$ space. We will try to mimic this approach in more general settings.

In the rest of this section, we will not use any assumptions on the trait space $(x)$ or the environmental space $(y)$. We will introduce a projection $M(u)$, playing the role of the ratio $R(u)$ used in the previous section, and a similar environmental projection $\Phi(\varphi)$ and consider the dynamics in the $(M, \Phi)$-plane. While $\partial M/\partial t$ and $\partial \Phi/\partial t$ can generally not be obtained from only the values of $M$ and $\varphi$, the key observation is that knowing the *signs* of these derivatives in an appropriate region is (almost) sufficient to deduce the strong fluctuations in $M$ and $\Phi$.

Let us state more precise hypotheses on the signs of $\partial M/\partial t$ and $\partial \Phi/\partial t$ that are sufficient to guarantee continual evolution. Here we focus first on stating hypotheses that are as simple as possible. Afterwards we will discuss examples leading to more realistic but also more complicated assumptions that are still sufficient to obtain continual evolution.

Suppose that we have found a function $M(u)$, taking values in $\mathbb{R}$, and that the environmental projection $\Phi$ also takes on values in $\mathbb{R}$. Without loss of generality we may also assume that both $M$ and $\Phi$ take on values in the closed interval $[0, 1]$. We assume that all rates of change, both ecologically and those due to mutations, vary continuously with $u$ and $\Phi$. We assume that there exists values $0 < \Phi_- < \Phi_+ < 1$ such that the rates of change *due to ecology only* satisfy the following feedback conditions:

- PF1 When $\Phi < \Phi_+$ we assume that $\frac{\partial M}{\partial t} > 0$ whenever $M$ is unequal to, but sufficiently close to 1, and similarly:
- PF2 When $\Phi > \Phi_-$ we assume that $\frac{\partial M}{\partial t} < 0$ whenever $M$ is unequal to, but sufficiently close to 0.
- NF1 When $\Phi \geq \Phi_+$ and $M < 1$ we assume that $\frac{\partial M}{\partial t} < 0$. When $M = 1$ and $\Phi \leq \Phi_+$ we assume that $\frac{\partial \Phi}{\partial t} > 0$, and similarly:
- NF2 When $\Phi \leq \Phi_-$ and $M > 0$ we assume that $\frac{\partial M}{\partial t} > 0$. When $M = 0$ and $\Phi \geq \Phi_-$ we assume that $\frac{\partial \Phi}{\partial t} < 0$.

Note that (PF2) and (NF2) are the symmetric equivalents of the conditions (PF1) and (NF1). Although it is not necessarily the case, we think of the extremal values $M \in \{0, 1\}$ as extinction of all but one phenotype. Therefore we assume that (still when considering only the effects due to ecology) $\partial M/\partial t = 0$ when $M \in \{0, 1\}$.

Next, we introduce mutations. The simple fact that mutations are possible naturally leads to the assumption that $\partial M/\partial t < 0$ when $M = 1$, and similarly that $\partial M/\partial t > 0$ when $M = 0$.

Finally, we introduce time scale assumptions. We assume that the rates of change $\partial \Phi/\partial t$ and the effect of mutations on $\partial M/\partial t$ are both arbitrarily small. In particular given a compact region of the $(M, \Phi)$-domain where the rate of change $\partial M/\partial t$ due to ecology is known to be non-zero, we may assume that the rate will still be non-zero (with the same sign) after introducing the effects due to mutations.

**Theorem 3.** *Suppose that the assumptions introduced above are satisfied. Then there exist initial values $(u, \Phi)$ for which $M(t)$ will fluctuate infinitely often between values arbitrarily close to both $0$ and $1$.*

*Proof.* **Step 1.** Start with an initial value $(M, \Phi)$ where $M$ is close to $1$ and $\Phi < \Phi_+$.
**Step 2.** Assumption (PF1) guarantees that $M$ stays close to $1$, while assumption (NF1) plus continuity of derivatives guarantee that $\Phi$ grows steadily, until the value of $\Phi$ gets sufficiently close to $\Phi_+$.
**Step 3.** When $\Phi$ gets sufficiently close to $\Phi_+$, $\Phi$ remains increasing, while assumption (NF1) plus the effects of mutations guarantee that $M$ moves away from the value $1$ by some definite amount.
**Step 4.** Once $M$ is strictly smaller than $1$ by some definite amount while $\Phi$ is still at least arbitrarily close to $\Phi_+$, it follows from the continuity assumption that the rate of change $\partial M/\partial t$ due to mutations is strictly negative. By our assumption on the time scales, the effect of mutations does not change this. Thus $M$ starts to decrease even further, and as long as $\Phi$ remains at least arbitrarily close to $\Phi_+$, reaches a value arbitrarily close to $0$ in some bounded time by compactness considerations, due to the fact that $M < 1$ by some definite amount. By the time scale assumption the value of $\Phi$ can change only arbitrarily little in this time interval, hence does not negate the decrease in $M$.
**Step 5.** We have ended up with a new initial condition where $M$ is arbitrarily close to $0$ and $\Phi$ is strictly larger than $\Phi_-$. By the symmetry of the assumptions the process repeats itself. □

To apply this proof of the general case to a specific system, there is a critical issue with the assumptions (NF1) and (NF2). The assumption in (NF1) and (NF2) that the ecological part of $\partial M/\partial t$ is strictly negative when $\Phi > \Phi_+$ is unrealistic: when $u$ consists of a single phenotype, one cannot expect changes in $M$ due to ecology. Therefore, we have to rely on mutations to be able to pass from $M$ close to $1$ to $M$ close to $0$. However, we want to maintain the assumption that mutations occur at a slower time scale than changes in $M$ due to ecology. An extra assumption is then needed to guarantee that in Step 4 of the proof the value of $\Phi$ cannot decrease enough to interfere with the decrease in $M$ and thereby to guarantee that, although (NF1) and (NF2) are not generally true, we are always in the region where they are true.

We will suggest two alternative assumptions. The first assumption, which is simplest, is to assume that the time scale at which $\Phi$ changes is still arbitrarily slower than the time scale at which mutations take place. This may or not be a realistic assumption, depending on the environmental factor $\Phi$ that the model considers. A second solution, which we will discuss in more detail below, is to assume that mutations from any trait to any other trait are possible, with uniform bounds on the ratios of different mutation rates. While it may not be immediately apparent, we will prove that this assumption is still sufficient to obtain continual evolution.

S1.3. **One evolving trait with a continuous range of phenotypes.** We will now present a more explicit setting in which we can prove continual evolution for a continuous range of traits $x \in \mathbb{R}$ and a single environmental factor $\varphi \in \mathbb{R}$, corresponding to a single point $y$. The interested reader will have no difficulty translating the results in this section to a discrete space $x$. In fact, all arguments literally hold true when the $L_1$ norm is replaced by the $\ell_1$ norm.

The argument closely follows the strategy presented previously: We will introduce a one-dimensional projection $M(u)$, and consider the dynamics in the $(M, \varphi)$-plane. While in the model it will not be possible to deduce the value of $\partial M/\partial t$ from only the values of $M$ and $\varphi$, the key observation is again that knowing the *sign* of $\partial M/\partial t$ in an appropriate region is (almost) sufficient to deduce the strong fluctuations in $M$ and $\varphi$.

Without loss of generality we consider a population density $u$ depending on traits $x \in [0, 1]$. As before, we will stipulate a fast positive feedback and a slow but dominating negative feedback caused by an environmental factor $\varphi$. We will assume that $\varphi$ is produced more by traits with larger $x$, and for fixed values of $u = (u_0, \dots, u_1)$ converges towards the average index $M$ given by

$$M := \frac{\|xu\|}{\|u\|},$$

where $\|\cdot\|$ represents the $L^1$-norm on the interval $[0, 1]$, e.g.

$$\|xu\| = \int_0^1 xu(x)\,dx.$$

Looking first only at the ecological factors of the dynamical system, ignoring for the moment the effects due to mutations, we assume that the densities of the individuals with trait $x$ change according to the logistic model

$$\frac{\partial u}{\partial t}(x) = u(x)\left(\mu_x(u, \varphi)\left(1 - \frac{\|u\|}{K}\right) - d\right),$$

where $K > 0$ is the carrying capacity, $d > 0$ the death rate and each growth rate $\mu_x$ is strictly positive, depends continuously on $x$, $u$ and $\varphi$, and we assume that for any given values of $u$ and $\varphi$ the dependence of $\mu$ on $x$ is either constant or strictly monotonic.

We further assume that the death rate is such that $\|\frac{\partial u}{\partial t}\| > 0$ whenever $\|u\|$ is sufficiently small, hence the total population size stays bounded from below. It is clear that the total population size automatically stays bounded from above by $K$.

When $\mu$ is constant the relative population sizes do not change, and it follows that $\frac{\partial M}{\partial t} = 0$. The monotonicity of $\mu$ will be used to determine the sign of $\frac{\partial M}{\partial t}$:

**Lemma 4.** *When $\mu$ is strictly increasing (resp. decreasing) in $x$, the rate $\frac{\partial M}{\partial t}$ is strictly positive (resp. negative).*

*Proof.* Let us assume that $\mu$ is strictly increasing, the argument is identical when $\mu$ is decreasing. We note that

$$\begin{aligned}\frac{\partial M}{\partial t} &= \frac{\partial}{\partial t}\left(\frac{\|xu\|}{\|u\|}\right)\\ &= \frac{\|x\frac{\partial u}{\partial t}(x)\| \cdot \|u\| - \|xu\| \cdot \|\frac{\partial u}{\partial t}(x)\|}{\|u\|^2}.\end{aligned}$$

Since we are currently only interested in the sign of $\frac{\partial M}{\partial t}$, we can drop the denominator and hence need to prove that

$$\frac{\|x\frac{\partial u}{\partial t}(x)\|}{\|xu(x)\|} > \frac{\|\frac{\partial u}{\partial t}(x)\|}{\|u\|}.$$

Plugging in the formula for $\frac{\partial u}{\partial t}(x)$, we note that the terms $(1 - \frac{\|u\|}{K})$ and $-d$, which are both independent of $x$, drop out of the quotients, and we are therefore left with showing that

$$\frac{\|x\mu(x) \cdot u(x)\|}{\|xu(x)\|} > \frac{\|\mu(x) \cdot u(x)\|}{\|u\|},$$

which is equivalent to

$$\frac{\|x\mu(x) \cdot u(x)\|}{\|\mu(x) \cdot u(x)\|} > \frac{\|xu(x)\|}{\|u\|}.$$

Since $\mu$ is assumed to be increasing in $x$ this inequality holds. □

We will assume that the rate of change of the environmental factor $\varphi$ can be written as

$$\frac{d\varphi}{dt} = \epsilon_e \cdot h(u, \varphi),$$

where the constant $\epsilon_e$ will later be assumed to be sufficiently small and $h$ is continuously differentiable. We make the following negative feedback assumptions, corresponding to the conditions with the same name discussed in section S1.2 discussing the general case:

(NF1) The function $h$ satisfies $h > 0$ when $\varphi < M$. Moreover, there exist $0 < \varphi_+ < 1$ such that for any non-zero $u$ the rate $\mu$ is decreasing in $x$ for $\varphi > \varphi_+$.

(NF2) The function $h$ satisfies $h < 0$ when $\varphi > M$. Moreover, there exist $0 < \varphi_- < 1$ such that for any non-zero $u$ the rate $\mu$ is increasing in $x$ for $\varphi < \varphi_-$.

A simple example of a function $h$ satisfying the first condition in each of these assumptions is $h(u) = M(u) - \varphi$. In general it may not be possible to express the function $h$ in terms of $M$ and not in $u$. These two assumption implies that for each value of $M$ there is a unique value $\varphi = \varphi(M)$ for which $h = 0$. The assumption that this unique value equals $M$ is merely a convenience, which by the implicit function theorem can be obtained by a change of coordinates whenever $\frac{\partial h}{\partial \varphi}(u, \varphi) \neq 0$ is satisfied for all $u$ and $\varphi = \varphi(M)$. We also note that necessarily $\varphi_+ > \varphi_-$.

For convenience we combine the two positive feedback assumptions (PF1) and (PF2) into the following assumption.

(PF) For each $\varphi$ there is a unique value $M_\varphi$ such that $\mu$ is strictly increasing at $(u, \varphi)$ whenever $M(u) > M_\varphi$, and strictly decreasing whenever $M(u) < M_\varphi$. We assume that the dependance of $M_\varphi$ on $\varphi$ is non-decreasing.

As a consequence of these feedback assumptions it follows that when $\mu$ is non-decreasing, it must remain so when $\varphi$ is decreased or when $M(u)$ is increased, and in the latter case must become strictly increasing. Similarly, when $\mu$ is non-increasing, it must remain so when $\varphi$ is increased or when $M(u)$ is decreased, in the latter case it must become strictly decreasing.

By continuity of $\mu$ it follows from (PF) that $\mu$ is constant when $M(u) = M_\varphi$. Note that we may redefine $\varphi_+$ as the smallest $\varphi$ for which $M_\varphi = 1$, and similarly $\varphi_-$ as the largest $\varphi$ for which $M_\varphi = 0$. Note also that we do not assume that $\mu$ is a function of $M$.

Let us now add mutations to the model:

$$\frac{du}{dt}(x) = u(x)\left(\mu(x)\left(1 - \frac{\|u\|}{K}\right) - d\right) + \epsilon_m \cdot \int_0^1 g_{\hat{x},x}(u)\, d\hat{x}.$$

Here $g_{\hat{x},x}$ represent mutations from $\hat{x}$ to $x$. We assume that $g$ is continuously differentiable in $x, \hat{x}$ and $u$, that $g_{x,\hat{x}} = -g_{\hat{x},x}$, and that $g_{\hat{x},x} = 0$ when $u(\hat{x}) = u(x) = 0$. We moreover assume that $g_{\hat{x},x} = 0$ is strictly increasing in $\hat{x}$, and thus strictly decreasing in $x$. In particular $\int_0^1 g_{\hat{x},x}(u)\, d\hat{x}$ is strictly positive when $u(x) = 0$ but $u \neq 0$, and mutations from any trait to any other trait are possible. We will later discuss alternative assumptions, making it possible to restrict the mutations that can occur.

**Lemma 5.** *Each $u(x)$ remains bounded from below by $C\epsilon_m$, where $C > 0$ can be chosen independently of $\epsilon_m$.*

*Proof.* Since the rate of change in $\|u\|$ due to mutations vanishes, our earlier assumptions on the death rate $d$ guarantee that $\|u\|$ remains bounded from above and below, i.e. $\|u\| = O(1)$. It follows that as $u(x) \to 0$:

$$\epsilon_m \cdot \int_0^1 g_{\hat{x},x}(u)\, d\hat{x} \geq O(\epsilon_m).$$

Thus for $u(x)$ small, the worst case scenario is that

$$\frac{\partial u}{\partial t}(x) \geq -c_1 \cdot u(x) + c_2 \cdot \epsilon_m,$$

for some uniform constants $c_1, c_2 > 0$ that are independent of $\epsilon_m$. It follows that

$$u(x) \geq \frac{c_2 \epsilon_m}{c_1} = C\epsilon_m.$$

□

**Theorem 6.** *For $\epsilon_e$ and $\epsilon_m$ sufficiently small there exist orbits $(u(t), \varphi(t))$ for which $M(t)$ alternates between values arbitrarily close to both 0 and 1 infinitely often.*

*Proof.* **Step 1.** Suppose that we start with initial values $u(t_0), \varphi(t_0)$ for which $M(t_0) := M(u(t_0)) > \varphi_+$, and for which $\mu$ is increasing in $x$. By our assumptions such initial values exist. It follows that $\varphi(t_0) < \varphi_+$, hence $\varphi(t_0) < M(t_0)$ and therefore $\varphi$ is increasing at time $t_0$.
**Step 2.** By choosing $\epsilon_m$ sufficiently small we can guarantee that $M(t)$ remains arbitrarily close to 1 until $\varphi > \varphi_+ - \delta$, where $\delta > 0$ can be chosen arbitrarily small. Note that $\varphi$ remains increasing while $\varphi \leq \varphi_+ - \delta$, hence at some time $t_1 > t_0$ we must have $M(t_1) = M_{\varphi(t_1)}$. We may assume that $t_1$ is the first time that this occurs, from which it follows that $M(t_1)$ must still be arbitrarily close to 1, while $\varphi(t_1)$ must be arbitrarily close to $\varphi_+$. In particular $\varphi$ is still increasing at time $t_1$, while $M(t)$ is decreasing due to mutations. As a consequence (PF) implies that $\mu$ becomes strictly decreasing, hence $M(t)$ will continue to decrease.
**Step 3.** Since $\varphi$ remains increasing and $M(t)$ remains decreasing, it follows that there is a smallest time $t_2 > t_1$ for which $M(t_2) = \varphi(t_2)$. We may assume that $t_2$ is the first time at which equality occurs, from which it follows that $\varphi$ has only increased between $t_1$ and $t_2$, and hence $\varphi(t_2) > \varphi_+ - \delta$.

We claim that at time $t_2$ we have that $u(0)$, the population density at $x = 0$, is bounded from below by a constant that is independent from the choice of $\epsilon_m$. To see this, note that since $\|u\|$ remains bounded, it follows that $u(x)$ for $x \in [0, 1-\xi]$ must remain comparable to $\epsilon_m$ for $t \in (t_0, t_1)$ for some $\xi$ that can be chosen arbitrarily small. It follows that the corresponding growth factors $\mu(x)\left(1 - \frac{\|u\|}{K}\right) -$

$d$ must remain strictly negative, with a uniform bound from above. It follows that populations $u(x)$ for $x \leq 1-\xi$ remain comparable to $\epsilon_m$, where Lemma 5 implies the estimate from below, and in particular the populations $u_x(t_1)$ for $x \leq 1-\xi$ are comparable to each other, with ratios independent of $\epsilon_m$. Recall that for $t \in (t_1, t_2)$ we noted that $\mu$ is increasing in $x$, and hence the growth factor $\mu_i\left(1-\frac{\|u\|}{K}\right)-d$ is largest for $x=0$. It follows that in the interval $[t_1, t_2]$ the population $u_0$ grows faster than any other population $u_i$, with a strictly larger exponential coefficient. At time $t_2$ the average $M(t)$ has decreased by an amount independent of $\epsilon_m$. Since the total population remains bounded away from 0 by assumption, it follows that the size of $u_0$ must have increased by an amount independent of $\epsilon_m$, thus obtaining the claim.

**Step 4.** By assuming that $\epsilon_m$ and $\epsilon_e$ are sufficiently small, it follows from (PF) and continuity of $\mu$ that $\mu$ will remain decreasing and $M(t)$ decreases below $\mu_-$, say at time $t_3$, and that the time interval $t_3 - t_2$ is bounded and independent of $\epsilon_m$ or $\epsilon_e$. Since $\epsilon_e$ is assumed to be small, it follows that $\varphi(t_2) \sim \varphi(t_1)$.
**Step 5.** We have ended up with assumptions on $M(t_3)$ and $\varphi(t_3)$ that are symmetrical to those on $M(t_0)$ and $\varphi(t_0)$. The symmetry of our assumptions implies that the process will repeat itself infinitely often, causing arbitrarily large fluctuations in $M(u)$. $\square$

If we drop the assumption that mutations from any strain to any other strain are possible, and replace it instead by the much weaker assumption that given any two traits there is a possible sequence of mutations from one to the other, Lemma 5 fails, and the above proof breaks down. We cannot guarantee that at time $t_2$ the trait $u(0)$ has increased to a definite size, independent of $\epsilon_m$, and as a result we cannot give a bound on the time interval $t_3 - t_2$.

This issue can be solved by assuming that the constant $\epsilon_e$ is sufficiently small, where the bound on $\epsilon_e$ may have to depend on the choice of $\epsilon_m$. Note the difference with the above statement, which holds whenever both $\epsilon_m$ and $\epsilon_e$ are sufficiently small. In practice the stronger assumption on $\epsilon_e$, which can imply that $\epsilon_e$ is much smaller than $\epsilon_m$, may or may not be desirable.

### S2. Examples: Rate equations, diversity index and supplementary figures

S2.1. **Shannon index.** The Shannon diversity index (H) is used in ecological research to describe species richness. Here we used the measure to describe phenotype richness within a species (we do not discuss whether bimodal phenotype distributions lead to different species). We calculate this diversity with the following formula:

$$H = -\sum_{1}^{n} p_i \ln p_i$$

Where $n$ is the number of phenotypes and $p_i$ the proportion of phenotype $i$ in the population.

S2.2. **One evolving trait with two phenotypes.** We have two examples for the case of a species with a single evolving trait with only two possible phenotype. In the first example we keep the total population constant and use $R$ as the ratio between the two subpopulations. We use the following equations:

$$\frac{dR}{dt} = R\left(\frac{0.011 + \frac{R}{1+R}}{k_A\left(1+\frac{\varphi}{k_B}\right)\left(1+\frac{0.011+\frac{R}{1+R}}{k_A}\right)} - 0.5\right) + \epsilon_M \frac{1-R}{1+R}$$
$$\frac{d\varphi}{dt} = \epsilon_E\left(\frac{R}{1+R} - \varphi\right)$$

$R$ and $\varphi$ are short for $R(t)$ and $\varphi(t)$ since they are time dependent. $R$ is the ratio of the two phenotypes, $A$ and $B$ ($R = \frac{A}{B}$), where $A$ is the phenotype interacting with the negative environmental factor $\varphi$. Figure S2 shows the result of time simulations of this example.

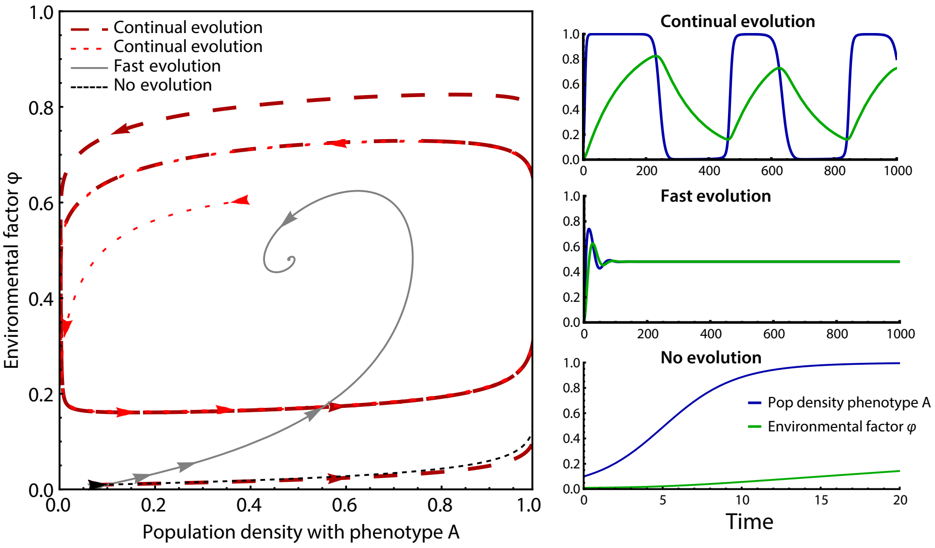


FIGURE S2. **Example with a single evolving trait with two phenotypes with a fixed population size** Behaviour of a system with two phenotypes with a fixed population size. A phase plane diagram (left) of population density with phenotype A and the environmental factor $\varphi$ shows the possible system behaviour depending on the relative timescales of population dynamics, the environment and mutations. The density of the subpopulation with phenotype B is 1 minus the subpopulation size with phenotype A. Slow environmental feedback and mutations leads to continual cyclic evolutionary dynamics, almost independent of initial conditions (right top). With fast environmental feedback and mutations an equilibrium is reached (right middle). No evolution (right bottom) leads to the extinction of one of the subpopulation (in this case the one with phenotype B). We used $k_A = 0.01$ and $k_B = 0.5$ throughout the figure. For the continual evolution we used $\epsilon_E = 0.008$ and $\epsilon_M = 0.0005$, for the fast evolution we used $\epsilon_E = 0.1$ and $\epsilon_M = 0.01$ and for no evolution we used $\epsilon_E = 0.01$ and $\epsilon_M = 0$

A second example, without the constant population size, is shown in Figure 5 in the main text and the equations for the time derivatives of the subpopulations with phenotypes $A$ and $B$ and the environmental factor $\varphi$ for that figure are:

$$\frac{dA}{dt} = A\left(\frac{(0.011+A)\left(1-\frac{A+B}{K}\right)}{k_A\left(1+\frac{\varphi}{k_B}\right)\left(1+\frac{0.011+A}{k_A}\right)} - d\right) + \epsilon_M(B-A)$$

$$\frac{dB}{dt} = B\left(0.5\left(1-\frac{A+B}{K}\right) - d\right) + \epsilon_M(A-B)$$

$$\frac{d\varphi}{dt} = \epsilon_E\left(A-\varphi\right)$$

$A$, $B$ and $\varphi$ are all time dependent and therefore short for $A(t)$, $B(t)$ and $\varphi(t)$. $k_A$ and $k_B$ are parameters describing the growth of $A$, $d$ is a death rate and $K$ the carrying capacity. The parameters used for Figure 5 are $k_A = 0.01$, $k_B = 0.5$, $d = 0.3$ and $K = 10$. For the continual evolution we used $\epsilon_E = 0.0005$ and $\epsilon_M = 0.00005$, for the fast evolution we used $\epsilon_E = 0.1$ and $\epsilon_M = 0.01$ and for no evolution we used $\epsilon_E = 0.01$ and $\epsilon_M = 0$.

S2.3. **One evolving trait with a range of phenotypes.** The equations used for Figure 3 in the main text are:

$$\frac{du_i}{dt} = u_i\left(\left[\frac{1}{2} + i\left(\frac{1}{3}(M-\frac{1}{2})) - \frac{2}{3}(\varphi-\frac{1}{2})\right)\right]\left(1-\frac{U}{K}\right) - d\right) + \epsilon_M\sum_{j\in I}\frac{1}{n}(u_j-u_i)e^{-10|i-j|}$$

$$\frac{d\varphi}{dt} = \epsilon_E\left(M-\varphi\right)$$

The collection of phenotypes $I$ consists of $n$ phenotypes with trait value $i$ and subpopulation density $u_i$. The total population size is $U = \sum_i u_i$. The average value of the trait in the population is $M = \frac{\sum_{i\in I} i\cdot u_i}{U}$. $\varphi$ is the level of the environmental factor, $K$ the carrying capacity and $d$ the death rate.

The rate with which mutations change the phenotype decreases exponentially with the difference between the phenotypes. This is to simulate that mutations with small effect are more prevalent than mutations with large effects.

We did a time simulation with different values of the rate of evolution ($\epsilon_M$) and the delay in the change of the environmental factor ($\epsilon_E$). For Figure 3 in the main text we used the following parameters: $K = 1$ and $d = 0.01$ throughout and for the continual dynamics $\epsilon_E = 10^{-4}$ and $\epsilon_M = 10^{-6}$, for the fast evolution $\epsilon_E = 0.1$ and $\epsilon_M = 0.01$ and when there is no evolution $\epsilon_E = 10^{-4}$ and $\epsilon_M = 0$. We assumed a subpopulation to go extinct if the subpopulation size was under 0.005.

S2.4. **Two evolving traits.** We simulated a system with two traits associated with two environmental factors for Figure 6 in the main text. The time derivatives of the subpopulations $u_i$, where $i = \{i_1, i_A\}$ is a vector of the two subpopulations, are:

$$\frac{du_i}{dt} = u_i\left(\frac{1}{2} + \frac{1}{2}\left[i_1\left(\frac{1}{3}(M_1-\frac{1}{2}) - \frac{2}{3}(\varphi_1-\frac{1}{2})\right) + i_A\left(\frac{1}{3}(M_A-\frac{1}{2}) - \frac{2}{3}(\varphi_A-\frac{1}{2})\right)\right]\left(1-\frac{U}{K}\right) - d\right) + \epsilon_M\left(\sum_{j\in I|j_1=i_1}(u_j-u_i)e^{-10|i_A-j_A|} + \sum_{j\in I|j_A=i_A}(u_j-u_i)e^{-10|i_1-j_1|}\right)$$

$$\frac{d\varphi_1}{dt} = \epsilon_E\left(M_1-\varphi_1\right)$$

$$\frac{d\varphi_A}{dt} = \frac{1}{4}\epsilon_E\left(M_A-\varphi_A\right)$$

The collection of phenotypes $I$ consists of $n$ phenotypes with trait value $\{i_1, i_A\}$ and subpopulation density $u_i$. The total population size is $U = \sum_i u_i$. The average value of the first trait in the population is $M_1 = \frac{\sum_{i\in I} i_1 \cdot u_i}{U}$ and of the second trait $M_A = \frac{\sum_{i\in I} i_A \cdot u_i}{U}$. $\varphi_1$ is the level of the environmental factor for the first trait, $\varphi_A$ the level of the environmental factor for the second trait, $K$ the carrying capacity and $d$ the death rate.

Since mutations are rare we ignore mutations in both traits at the same time and the rate of mutations in one trait from one phenotype to the other decreases exponentially with increasing difference between the phenotype.

We used the parameters $d = 0.01$, $K = 1$, $\epsilon_E = 0.0001$ and $\epsilon_M = 0.0005$ for Figure 6 in the main text.

S2.5. **Literature models.** We used the competitor-competitor model from Khibnik and Kondrashov[2] where they show RQ dynamics (Fig. 2 in Khibnik and Kondrashov[2]). We changed the model to allow for a polymorphic population, which leads to the following equations:

$$\frac{dx_i}{dt} = x_i\left(r_{1,i} - r_2\sum_i x_i - \sum_j r_{3,i,j}y_j\right) + \epsilon_M\sum_{k\in I}\frac{1}{n}(x_k - x_i)e^{-10|i-k|}$$

$$\frac{dy_j}{dt} = y_j\epsilon_E\left(r_{4,j} - r_5\sum_j y_j - \sum_i r_{6,i,j}x_i\right) + \epsilon_M\sum_{k\in J}\frac{1}{n}(y_k - y_j)e^{-10|j-k|}$$

The parameters are given in Khibnik and Kondrashov[2] Eqs 6 and Fig. 2. $\epsilon_E$ is set to 1 in Fig. S3B and to 0.1 in Fig. S3C. The trait values ($i$ and $j$) range from 0.5 to 1.5 and we simulated 100 different phenotypes per species within this range.

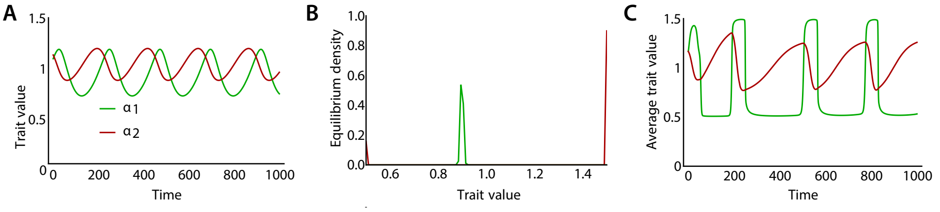


FIGURE S3. **RQ dynamics disappears when adding polymorphism to a population.** We have adapted a competitor-competitor model from[2] with originally an adaptive dynamics approach that previously showed Red Queen Dynamics to an instance with our assumptions (the possibility of a polymorphic trait distribution and larger mutations). **A** The competitor-competitor model of Khibnik and Kondrashov[2] shows the evolvable trait ($\alpha$) of the two competitors following each other in a RQ manner (figure replicated from Figure 2 in Khibnik and Kondrashov[2]). **B** When we allow for a polymorphic population the system tends to a bimodal distribution for the trait $\alpha_2$, and the RQ dynamics disappear. **C** Using the results from this paper that a slow negative feedback allows for RQ dynamics, we change the timescales and make the feedback (competitor species in this example) 10 times slower. The RQ dynamics are retrieved.

### S3. SUPPLEMENTARY TABLES

TABLE S1. Additional examples of negative and positive feedbacks.

| **Examples of slow negative feedbacks** |
|---|
| • a herbivore that changes the vegetation which then becomes less suitable for the herbivore |
| • micro-organisms that produce a compound that inhibits them but is diluted in a large volume (to achieve the feedback being slow) |
| • a species with a complex life cycle with a habitat shift (with an evolvable component), if a crowded habitat has a delayed effect on the habitat quality whole ecosystem that leads to a negative feedback (e.g. with intransitive cycles as in Bonachela et al. [1]) |
| • a feedback could also be caused by human intervention, such as flu vaccinations—common viruses will be vaccinated against during the next season |
| **Examples of fast positive feedbacks** |
| • In the case of a habitat shift in the life cycle, similar habitat shift will share the second habitat with more individuals, providing mates or protection |

MEIKE T. WORTEL AND NILS CHRISTIAN STENSETH: CENTRE FOR ECOLOGICAL AND EVOLUTIONARY SYNTHESIS (CEES), DEPARTMENT OF BIOSCIENCES, UNIVERSITY OF OSLO, OSLO, NORWAY

HAN PETERS: KORTEWEG DE VRIES INSTITUTE FOR MATHEMATICS, UNIVERSITY OF AMSTERDAM, AMSTERDAM, THE NETHERLANDS

PRESENT ADDRESS MEIKE T. WORTEL: INSTITITE FOR BIODIVERSITY AND ECOSYSTEM DYNAMICS, UNIVERSITY OF AMSTERDAM, AMSTERDAM, THE NETHERLANDS

JUAN A. BONACHELA: DEPARTMENT OF ECOLOGY, EVOLUTION, AND NATURAL RESOURCES, RUTGERS UNIVERSITY, NEW BRUNSWICK, U.S.A.
